# Adaptive RAxML-NG: Accelerating Phylogenetic inference under Maximum Likelihood using dataset difficulty

**DOI:** 10.1101/2023.05.15.540873

**Authors:** Anastasis Togkousidis, Alexey M. Kozlov, Julia Haag, Dimitri Höhler, Alexandros Stamatakis

## Abstract

**Motivation:** Phylogenetic inferences under the Maximum-Likelihood (ML) criterion deploy heuristic tree search strategies to explore the vast search space. Depending on the input dataset, searches from different starting trees might all converge to a single tree topology. Often, though, distinct searches infer multiple topologies with large log-likelihood score differences or yield topologically highly distinct, yet almost equally likely, trees. Recently, Haag *et al*. introduced an approach to quantify, and implemented machine learning methods to predict, the difficulty of an MSA with respect to phylogenetic inference. Easy MSAs exhibit a single likelihood peak on their likelihood surface, associated with a single tree topology to which most, if not all, independent searches rapidly converge. However, as difficulty increases, multiple locally optimal likelihood peaks emerge, yet from highly distinct topologies.

**Results:** To this end, we introduce and implement an adaptive tree search heuristic in RAxML-NG, which modifies the thoroughness of the tree search strategy as a function of the predicted difficulty. Our adaptive strategy is based upon three observations. First, on easy datasets, searches converge rapidly and can hence be terminated at an earlier stage. Second, over-analyzing difficult datasets is hopeless and, thus, it suffices to quickly infer only one of the numerous almost equally likely topologies, to reduce overall execution time. Third, more extensive searches are justified and required on datasets with intermediate difficulty. While the likelihood surface exhibits multiple locally optimal peaks in this case, a small proportion of them is significantly better. Our experimental results for the adaptive heuristic on 9, 515 empirical and 5, 000 simulated datasets with varying difficulty exhibit substantial speedups, especially on easy and difficult datasets (53% of total MSAs), where we observe average speedups of more than 10x. Further, approximately 94% of the inferred trees using the adaptive strategy are statistically indistinguishable from the trees inferred under the standard strategy (RAxML-NG).

**Availability:** GNU GPL at https://github.com/togkousa/raxml-ng/tree/adaptive.

**Supplementary Material:** Available

## 1 INTRODUCTION

Phylogenetic tree inference addresses the problem of finding the binary tree that best explains the sequence data, typically given in the form of a multiple sequence alignment (MSA). To infer trees, various techniques and methods have been developed, such as distance-based approaches (e.g., Neighbor Joining (Saitou and Nei, 1987)), Maximum Parsimony (MP) (Fitch, 1971), Maximum Likelihood (ML) (Felsenstein, 1981), and Bayesian Inference (BI) methods (Yang and Rannala, 1997; Mau *et al*., 1999). ML and BI methods rely on the phylogenetic likelihood function, which implements an explicit statistical model of sequence evolution. Computing the likelihood score of a given, single candidate tree already constitutes a computationally expensive task. ML and BI analyses are time- and resource-intensive because hundreds of thousands of likelihood computations are performed on a large number of distinct tree topologies. Here, we focus on developing adaptive heuristics for ML based phylogenetic inference. Analogous techniques could be developed for MP, BI, and potentially also NJ.

In addition to the computational burden associated with the likelihood function itself, ML inference is known to be an NP-hard problem (Roch, 2006) as the number of possible topologies grows super-exponentially with the number of sequences. Since the brute-force evaluation of all possible tree topologies is impossible, there exists a plethora of inference tools which deploy different heuristics to find a tree with a “good” likelihood score. In fact, heuristics are involved in all distinct phases of the ML tree search, including: (*i*) the construction of the starting tree(s) upon which one initiates the search, (*ii*) the underlying strategy for topological alterations, such as *Nearest Neighbor Interchange (NNI)* or *Subtree Prune and Regraft (SPR)* moves, in order to efficiently search the vast tree space, (*iii*) the optimization techniques applied to continuous parameters of the evolutionary model (e.g., branch lengths and substitution rates between states), (*iv*) the stopping criteria to terminate the tree search (St. John, 2016). Some of the most widely used tools for ML tree inference are RAxML (Stamatakis, 2014) and RAxML-NG (Kozlov *et al*., 2019), IQ-TREE 2 (Minh *et al*., 2020), FastTree 2 (Price *et al*., 2010) and PhyML (Guindon *et al*., 2010).

Heuristics, evidently, do not guarantee that one will find the globally optimal tree. Empirical observations (Morrison, 2007; Stamatakis, 2011; Morel *et al*., 2020) suggest that, on certain datasets, independent ML tree searches converge to a single - or topologically highly similar - tree(s), while on other datasets, they yield multiple topologically highly distinct trees with almost identical likelihood scores. In the latter case, standard phylogenetic significance tests, for example the *tree topology tests* implemented in IQ-TREE 2 (Naser-Khdour *et al*., 2019), usually report no statistically significant difference among the majority of the inferred, highly-contradicting, topologies. Thus, “easy” MSAs exhibit a single, well-distinguishable, globally optimal peak on their likelihood surface, associated with a single tree topology. In contrast, “difficult” MSAs exhibit a rugged likelihood surface, with multiple locally optimal and statistically indistinguishable peaks emerging from contradicting topologies.

This MSA behaviour is quantified by the Pythia tool (Haag *et al*., 2022b). In the corresponding paper, Haag *et al*. introduce the concept of MSA analysis *difficulty*. The difficulty is a real number between 0.0 to 1.0 that reflects the degree of ruggedness on the respective likelihood surface. Easy MSAs, with a difficulty score close to 0.0, exhibit a single globally optimal peak on their likelihood surface, while difficult ones, with a score closer to 1.0, exhibit multiple locally optimal peaks. The difficulty score does not only capture these two extreme cases, but also the entire spectrum of intermediate cases. Therefore, one can roughly classify MSAs into easy-, intermidate-, and hard-to-analyze. The Pythia tool uses machine learning methods to predict the difficulty given the MSA *prior* to phylogenetic analysis (Section 2.2).

Here, we introduce an adaptive ML tree search heuristic and implement it in RAxML-NG, based on Pythia’s difficulty prediction for the respective input MSA. Our new *adaptive RAxML-NG* tool modifies the thoroughness of the tree search strategy, as well as additional heuristic search parameters (e.g., the number of distinct starting trees or the maximum subtree re-insertion radius of SLOW-SPR moves), as a function of the predicted difficulty. We further introduce two new mechanisms for faster and more efficient exploration, that is, NNI rounds and the 1% ML convergence interval to terminate the first more superficial phase of topological moves early. We provide a detailed description of the new algorithm and the underlying mechanisms in Sections 2 and 3. Our experimental results (Section 4) yield substantial speedups, on 9, 515 empirical and 5, 000 simulated datasets, with varying difficulty under a representative difficulty distribution. The average speedup ranges between 1.8x for intermediate datasets and up to 16x for easy and difficult datasets. The overall accumulated speedup achieved over all datasets is approximately 3.4x. Further, in about 94% of the cases, the output ML trees from standard and adaptive RAxML-NG are statistically indistinguishable. All MSAs we used for our experiments, are available for download at https://cme.h-its.org/exelixis/material/raxml_adaptive_data.tar.gz.

Our main goal is to deploy Pythia difficulty scores for informing adaptive tree search heuistics under ML. To this end, we compare the standard and adaptive RAxML-NG versions in terms of topological accuracy and ML score of the respective output trees. We conduct experiments with a sufficiently large number of empirical and simulated datasets. Furthermore, our datasets are also representative of the difficulty distributions in phylogenetic data repositories such as TreeBASE (Piel *et al*., 2009) or RAxML Grove (Höhler *et al*., 2021). To simplify the experimental setup, we executed RAxML-NG and adaptive RAxML-NG in sequential execution mode only. In Section 5 we outline directions of future work including potential parallelization strategies for adaptive RAxML-NG.

## 2 RELATED WORK

### 2.1 ML tree search heuristics

The likelihood function has both, discrete and continuous parameters. The tree topology and the specific model of sequence evolution constitute discrete parameters, while the branch lengths of the input tree, the substitution rates between the characters/states, the equilibrium frequencies, and the *α* shape parameter of the Γ model of rate heterogeneity are continuous parameters (Yang, 2014). RAxML-NG optimizes the continuous parameters, with respect to the likelihood score, using the Newton-Raphson method for the branch lengths and other numerical optimization routines for the remaining parameters. It optimizes all continuous parameters iteratively, repeating the process for multiple rounds until either a numerical threshold is reached or no further changes are applied to the parameter values. ML inference tools typically split the optimization of continuous parameters into the *Branch-Length Optimization (BLO)* and the *Model-Parameter Optimization (MPO)* routines. The latter refers to any other continuous parameter except for branch lengths. Both, standard and adaptive RAxML-NG use the exact same framework to optimize continuous parameters.

Standard RAxML-NG exclusively explores the vast tree space via SPR moves (Figure 1a). The tree search heuristic is based on the greedy hill-climbing algorithm previously introduced in RAxML (Stamatakis, 2014), with minor modifications (Kozlov, 2018). Initially, the algorithm stores pointers to all inner nodes of the initial comprehensive tree topology (starting tree). For every pointer in the list, RAxML-NG prunes the corresponding subtree from the, thus far, best-scoring tree, and evaluates its possible re-insertions into neighboring branches, up to a certain maximum distance (reinsertion radius) from the pruning branch. The move that yields the highest likelihood improvement, if such a move exists, is accepted and the algorithm proceeds to the next element in the node list. When SPR moves for all pointers/nodes have been evaluated, and therefore all corresponding subtrees have been pruned and regrafted once, the SPR round is completed. There are two types of SPR rounds in *adaptive* RAxML-NG. During *Fast SPR rounds*, adaptive RAxML-NG evaluates each alternative tree topology that results from an SPR move, using the existing branch lengths, while in *Slow SPR rounds*, the lengths of the three adjacent branches around the insertion node are re-optimized. Further, in Slow SPR rounds the 20 top-scoring topologies are stored in a list *BT*. At the end of the round, all trees in *BT* undergo a full BLO and the best tree is used as initial tree for the next SPR round. The distinction between Fast and Slow SPR rounds is a feature of both RAxML and RAxML-NG. The implementation of Fast and Slow SPR rounds, differs among standard and adaptive RAxML-NG versions. However, the fundamental difference, that is the optimization of the adjacent branches around the insertion node, remains the same.

**Fig. 1.**
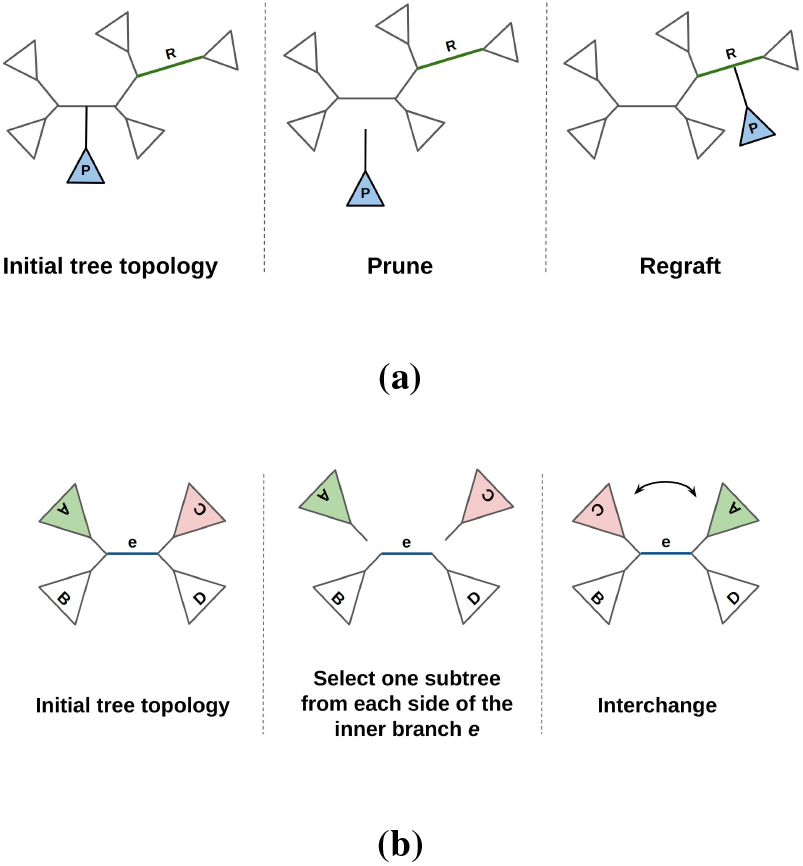
**(a)**An SPR move example. The subtree *P* is pruned from the initial comprehensive tree and regrafted into branch *R*. **(b)** An example of an NNI move around an inner branch *e*. Two subtrees, one from each side of the inner branch, are selected, pruned and interchanged. The NNI move is a special case of the SPR move.

In addition to SPR moves, our adaptive RAxML-NG vesion also uses NNI moves (Figure 1b) for tree space exploration. While NNI-based heuristics are more prone to becoming stuck in local optima, the main motivation for including NNI moves is that they might be sufficient to attain trees with good scores for easy datasets with a low difficulty score. The NNI algorithm is analogous to the *NNI round* implementation in RAxML v8.2 (-f J option). It starts from an inner branch and compares the three alternative NNI topologies in terms of their likelihood score. For computing the likelihood, adaptive RAxML-NG optimizes the five branches which are most affected by an NNI move. Once an NNI evaluation has been executed, the algorithm accepts the topology with the highest likelihood score and proceeds to the adjacent inner branches. When all inner branches have been visited once, the algorithm returns to the initial inner branch and repeats the process for multiple iterations, until it reaches an NNI-optimal tree (i.e., a tree where the application of any additional NNI move will not further improve the likelihood).

### 2.2 Difficulty Prediction

Pythia (developed in our research group) is a Gradient Boosted Tree Regressor^1^ trained to predict the difficulty of a phylogenetic analysis of an MSA *prior* to initiating ML-based tree inferences. Initially, we defined and calculated the inference difficulty of empirical MSAs by conducting 100 ML tree searches on each dataset using RAxML-NG. For each dataset, we extracted the *plausible tree set (PTS)*, that is, the inferred ML trees that are not significantly worse than the best-scoring ML tree under *any* statistical significance test implemented in IQ-TREE 2. Next, we calculated the Robinson–Foulds (RF) distances (Robinson and Foulds, 1981) between all pairs of trees in the ML tree set and in the PTS. The definition of the difficulty score is based on the proportion of plausible trees, the number of unique tree topologies in the ML tree set as well as in the PTS, and the average relative RF-distances among trees in the PTS.

Next, we trained and tested Pythia using the calculated difficulty labels. We selected eight features to represent each MSA as an eight-dimensional data point. Six of them are simple and fast-to-compute as they rely on MSA attributes (e.g., the proportion of gaps or the Bollback Multinomial metric (Bollback, 2002)). Pythia calculates the two remaining features by conducting 100 MP searches, which are orders of magnitude faster to compute than ML searches. The two features we extract from these 100 MP trees are the proportion of unique tree topologies in the MP tree set, and the average relative RF-distances between all pairs of MP trees. The results indicate high prediction accuracy with a mean absolute prediction error (MAE) of 0.09 and a mean absolute percentage error (MAPE) of 2.9% for Pythia. Note that, difficulty prediction is on average five times faster than a single ML tree inference. More details about the definition of the difficulty score can be found in the Supplementary Material and the original Pythia publication (Haag *et al*., 2022b).

### 2.3 Heuristics, difficulty, and phylogenetic signal

Properly evaluating and comparing different heuristics/tools constitutes a challenging task when introducing novel heuristics. According to a recent comparative study (Höhler *et al*., 2022) conducted in our research group, there is no standard set of benchmark data for assessing/comparing different tools. Most performance and accuracy studies, typically use ad hoc benchmark dataset collections. These collections sometimes exhibit specific properties that might even yield contradictory results (see examples further below). In this section we examine some major issues reported in the preprint by Höhler *et al*., in relation to previous studies as well. We outline the connection between the difficulty score of an MSA and the respective convergence speed of the ML heuristic, as well as the accuracy and robustness of the result.

FastTree 2 is one of the fastest and most widely used tools for ML inference. The tool applies a combination of “linear SPR” moves under MP and ML-NNI moves thereby achieving linear run time complexity with respect to the number of taxa (*O*(*n*)). The results in the FastTree 2 paper (Price *et al*., 2010) indicate that the tool is 100 −1, 000 times faster than PhyML and RAxML (version 7.2.1), but the trees inferred by the latter tools are substantially more accurate, due to the thoroughness of their tree search heuristics. This rather discouraging result concerning the inference quality of FastTree 2 is also confirmed by an independent study (Zhou *et al*., 2017) that also includes IQ-TREE (Nguyen *et al*., 2014). The authors found that RAxML, IQ-TREE, and PhyML perform similarly in terms of topological accuracy on single-gene datasets, while FastTree 2 performs substantially worse. In contrast to the aforementioned studies, the results from a third comparative study between RAxML (version 7.2.6) and FastTree 2 only (Liu *et al*., 2011), show that although RAxML generally infers topologically more accurate trees than FastTree 2, the differences diminish as the sites-over-taxa ratio decreases. A similar observation was also made by Höhler *et al*.^2^

Finally, the results from a recent study we conducted on SARS-CoV-2 data (Morel *et al*., 2020) should also be mentioned. Due to the comparatively low nucleotide substitution rate of SARS-CoV-2 (van Dorp *et al*., 2020), the four MSAs analyzed are characterized by a high proportion of invariable sites, and the patterns-over-taxa ratio is equal to or less than 1. We conducted 100 ML tree searches using RAxML-NG and generated the PTS. In all four MSAs, the independent searches yielded 100 distinct tree topologies, and the PTS comprised more than 70% of the 100 ML trees. The average relative RF-distance between all pairs of ML trees was approximately 0.78, implying topologically highly distinct trees.

The inconsistent conclusions in the aforementioned studies can potentially be explained by reflecting on the concept of the difficulty score. Starting from the latter study, the 100 output ML trees with contradicting topologies imply an extremely rugged likelihood surface that exhibits a plethora of local optima. We confirm this hypothesis by calculating the difficulty score of the full SARS-CoV-2 dataset; the score is 0.84 and the dataset is close to hopeless for analyzing. In other words, it will not yield a single strictly bifurcating tree, that is, a clear peak. Additional indications for a high degree of difficulty could also be the sites-over-taxa (SoT), and patterns-over-taxa (PoT) ratios of an MSA, which are also used by Pythia as prediction features. First, Rosenberg and Kumar (Rosenberg and Kumar, 2001) concluded that, the higher the SoT ratio of a dataset, the stronger the phylogenetic signal and the higher the accuracy of the result will be. This provides a sufficient explanation for the results of the aforementioned study by Liu *et al*. (2011). As the number of taxa grows, the SoT ratio of the MSA decreases and the difficulty increases. Hence, both RAxML and FastTree 2 infer (potentially) distinct locally optimal trees with similar likelihood scores, an observation also made by Höhler *et al*. Further, Figure 1 in Höhler *et al*. establishes a strong negative correlation between the PoT ratio and the difficulty of empirical MSAs, showing that the difficulty decreases substantially when the PoT ratio exceeds 10. This provides an additional explanation for the high difficulty of the SARS-CoV-2 dataset. In the same study, RAxML-NG and IQ-TREE 2 perform substantially better than FastTree 2 in terms of topological accuracy for empirical datasets with difficulties ranging between 0.2 and 0.6 (easy-to-intermediate) which is in agreement with the results of the two other studies (Price *et al*., 2010; Zhou *et al*., 2017). For easy datasets, though, with a difficulty score below 0.2, FastTree 2 performs similarly in terms of likelihood score and topological accuracy.

## 3 ALGORITHM

We begin by dividing datasets into three classes: (*a*) *easy* datasets with a difficulty score below 0.3, (*b*) *intermediate* datasets with a difficulty between 0.3 and 0.7, and (*c*) *difficult* datasets with a difficulty above 0.7. Our adaptive RAxML-NG tree search heuristic is based upon three observations derived from the discussion in Sections 2.2 and 2.3.

1. The majority of independent ML tree searches on easy MSAs converges to a single, or topologically highly similar, tree(s). Moreover, fast and thorough heuristics (FastTree 2 and standard RAxML-NG) perform similarly in terms of topological accuracy and likelihood score, especially when the difficulty score is close to 0.
2. Independent ML tree searches on difficult MSAs yield topologically highly distinct trees, with most of them being equally likely and, therefore, statistically indistinguishable. Fast and thorough heuristics perform similarly in this difficulty range, albeit only in terms of likelihood score. Overanalyzing such datasets is hopeless and, thus, it suffices to quickly infer only a few out of the many almost equally likely topologies, to reduce overall execution time.
3. On datasets of intermediate difficulty, thorough heuristics yield significantly (in the statistical sense) better ML trees than superficial ones. This is because faster heuristics are more prone to become stuck in local optima and, therefore, yield sub-optimal trees. In general, intermediate MSAs exhibit fewer peaks on their likelihood surface than the difficult ones. While the corresponding likelihood surfaces exhibit multiple peaks on intermediate datasets, only a small proportion of these peaks is significantly better. Thus, more extensive search strategies yield significantly better results.

We modify the number of independent ML tree searches conducted by RAxML-NG^3^ and the Slow SPR radius parameter - which directly determines the thoroughness of the tree search heuristic - according to the difficulty of the MSA. The functions to determine the number of random/MP starting trees depending on the respective difficulty in adaptive RAxML-NG are shown in Figure 2a. Both functions are Gaussian curves with a peak of 10 (trees) when the difficulty score is 0.5. We intentionally set a wider curve for the number of MP starting trees, such that the heuristic uses more MP than random starting trees on easy and difficult datasets. This is due to an observation on the difficult SARS-CoV-2 dataset. We observed that tree searches initiated on parsimony starting trees consistently yielded phylogenies with substantially better log-likelihood scores (Morel *et al*., 2020). In addition, Figure 2b shows the function to determine the Slow SPR radius depending on the difficulty. We use a triangle function, starting from a radius of 5 when the difficulty score is either 0 or 1 and a peak of 30 for a difficulty of 0.5. The numbers for random/MP starting trees and the Slow SPR radius parameter are then set as the integer part (floor) of the values from the corresponding functions.

**Fig. 2.**
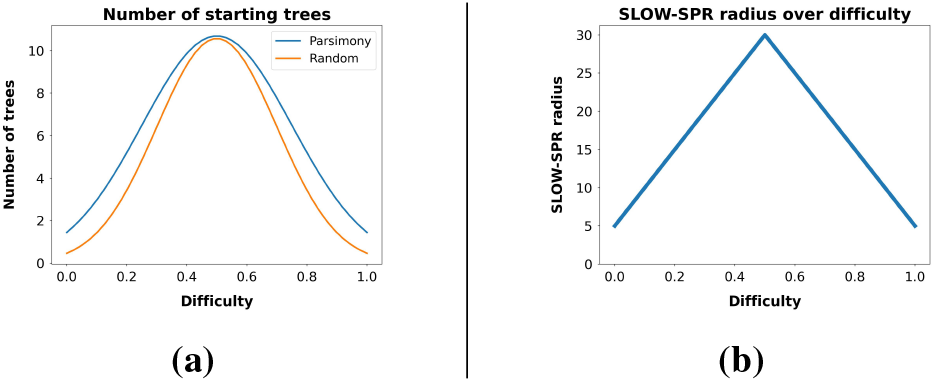
**(a)**Number of random/MP starting trees over difficulty **(b)** SLOW-SPR radius parameter over difficulty.

Standard RAxML-NG exclusively searches the tree space via SPR moves. Generally, the advantage of using NNI instead of SPR moves for tree searches, is that the time complexity of the heuristic is reduced from *O*(*n*^2^) to *O*(*n*), where *n* is the number of taxa (Heath and Ramakrishnan, 2010). On the other hand, NNI-based heuristics are more likely to become stuck in local optima and therefore infer sub-optimal trees. We trade speed for accuracy by deploying a combination of SPR and NNI moves. In our adaptive heuristic, every SPR Round (either Fast or Slow) is followed by an NNI round (see Section 2.1). The process is divided into two stages. During the first stage, Fast SPR rounds alternate with NNI rounds, until either the RF distance between two consecutive tree topologies is zero^4^ or the likelihood score improvement is below a user-defined threshold (Haag *et al*., 2022a). In the second stage, Slow SPR rounds alternate with NNI rounds until the likelihood score improvement threshold is reached again. Note that the algorithm optimizes the branch lengths (BLO) and model parameters (MPO) of the starting tree before entering the first stage. At the end of each stage it only re-optimizes the model parameters, since local and full BLOs are already part of the SPR and NNI rounds.

Finally, we introduce a new criterion for early termination of the first stage of the heuristic. We use the log-likelihood score of the best ML tree inferred so far by the already completed tree inferences, as a reference score to define a 1% ML convergence interval. We assume that the first stage of the tree search has converged when the log-likelihood score of the current tree is less than 1% worse than this reference score. We observed that alternating between SPR and NNI moves yields higher convergence speed, that is log-likelihood improvement over execution time, than standard RAxML-NG which exclusively relies on SPR rounds. For easy and difficult MSAs, adaptive RAxML-NG begins with an NNI round followed by MPO, as the probability to already converge when only using these two routines in those difficulty score ranges is comparatively high. In fact, around 70% of tree searches conducted on easy and difficult empirical/simulated MSAs converged by only applying an NNI round, followed by MPO. We provide the workflow of the adaptive RAxML-NG heuristic in Figure 3; the full pseudocode is available in the Supplementary Material.

**Fig. 3.**
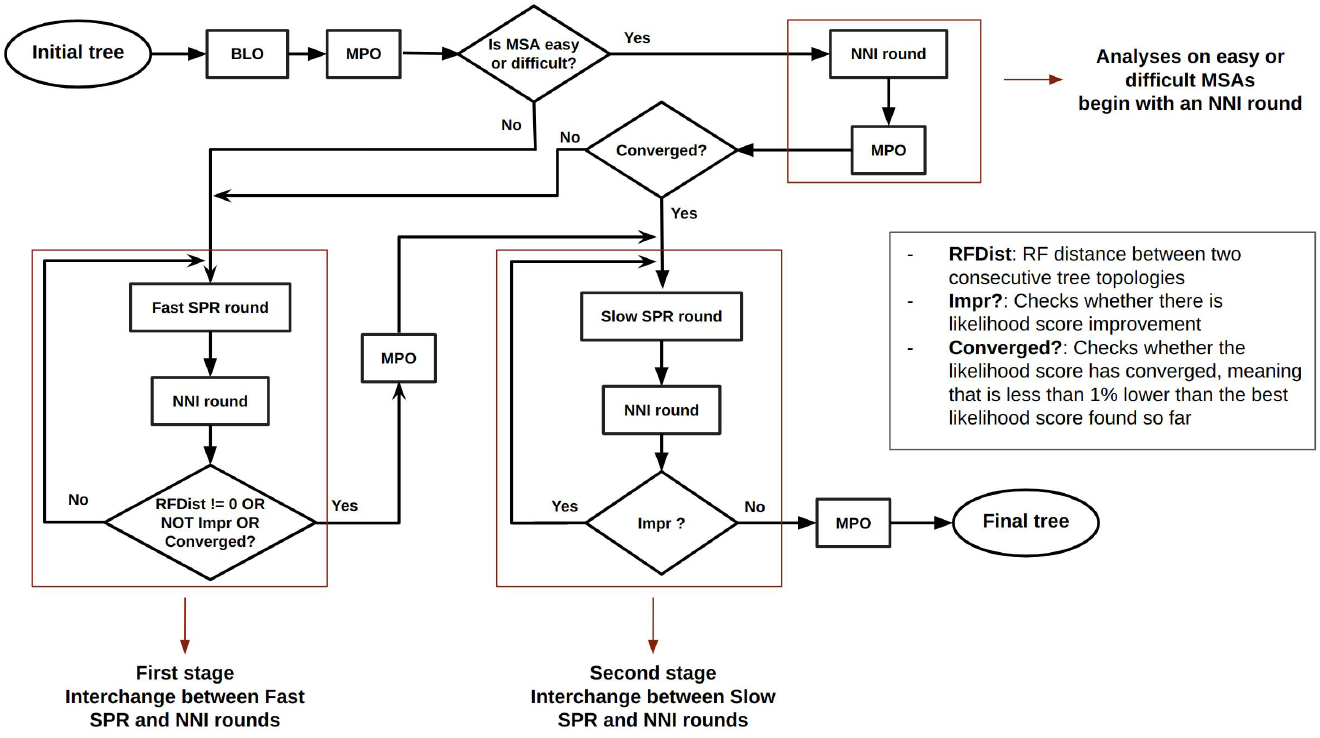
Workflow of a single adaptive RAxML-NG tree inference. Standard RAxML-NG, on the other hand, conducts three stages of SPR rounds. The first stage comprises SPR rounds “on fast mode”. The second and third stages are series of Fast and Slow SPR rounds respectively. The sole criterion to terminate a stage in standard RAxML-NG is the log-likelihood score improvement. Further, Fast and Slow SPR rounds differ between the standard and adaptive versions of RAxML-NG, although their basic principles remain the same (Kozlov, 2018).

## 4 RESULTS

We conducted experiments on 10, 389 empirical MSAs from TreeBASE (Piel *et al*., 2009) and 5, 000 simulated MSAs with varying difficulty. Regarding the simulated MSAs, we sampled 4, 500 of the already simulated DNA datasets used in the preprint by Höhler *et al*., and also generated an additional 500 simulated amino-acid datasets (see Supplement for details). We will refer to these MSAs as RAxML Grove simulated (RGS) datasets.

For each empirical and simulated MSA, we executed both, the standard and adaptive RAxML-NG versions in sequential mode. Due to the large number of MSAs, we set an execution time limit of 24 hours for standard and adaptive RAxML-NG. When tree searches with both RAxML-NG versions terminate within this pre-specified time interval, the experiment for the corresponding MSA is considered successful and we continue with further evaluations/comparisons. Otherwise, the dataset is discarded from downstream analysis.

We compare the two RAxML-NG versions based on: (*i*) the *log-likelihood score (LH)* of the output trees, (*ii*) the relative RF distance between them, (*iii*) the results from IQ-TREE 2 significance tests, and (*iv*) execution times. Regarding the significance tests, we consider the two output ML trees to be statistically indistinguishable, and therefore none of them to be significantly worse, if the corresponding RAxML-NG/adaptive RAxML-NG tree pair passes *all* seven statistical tests implemented in IQ-TREE 2. This is very conservative, but circumvents the discussion about the appropriateness of distinct statistical significance tests. In this case we assign the label PASSED to the pair. Otherwise, we assign the label FAILED. There are also some cases in which the execution of IQ-TREE 2 failed, due to unfavorable MSA properties (duplicated sequences or sequences only comprising gaps) or due to some invalid characters in taxon names provoking errors in IQ-TREE 2 when importing the trees inferred by RAxML-NG^5^. Those datasets were also discarded.

Due to this filtering, we present results for 9, 515 empirical and 5, 000 simulated MSAs, in which the execution of standard/adaptive RAxML-NG versions (within 24 hours of run time) and IQ-TREE 2 significance tests were successful. All simulated MSAs successfully passed all filtering steps. Figure 4 illustrates the distributions (density plots) of empirical/simulated datasets over ten difficulty intervals. We observe that the proportion of datasets with a difficulty score above 0.9 is comparatively low. This is also an observation made by Höhler *et al*. on TreeBASE and RGS datasets, and is associated with the definition of the difficulty score per se. More information regarding the datasets used can be found in the Supplementary Material.

**Fig. 4.**
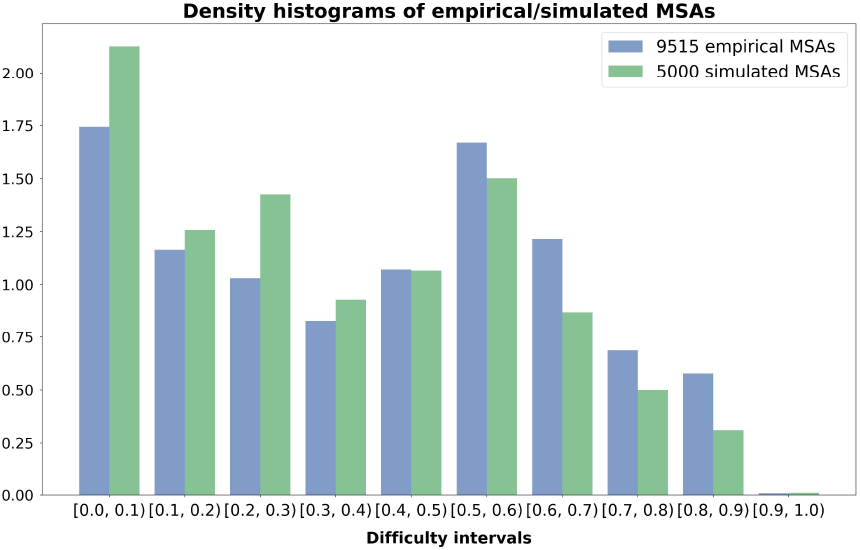
Density histograms showing the distribution of empirical/simulated MSAs on 10 difficulty intervals.

We ran our experiments on the Haswell/URZ Cluster, located at the Computing Center of the University of Heidelberg. It consists of 224 nodes with Intel Haswell CPUs (E5-2630v3 running at 2.40GHz). Each node contains 2 CPUs and each CPU has 8 physical cores. The operating system is CentOS Linux 7 (Core).

### 4.1 Comparing the log-likelihood scores

For the sake of simplicity, for a given MSA input, we will refer to the output ML tree inferred by standard RAxML-NG as the *standard tree*, and to the tree inferred by adaptive RAxML-NG as the *adaptive tree*. We define the absolute log-likelihood difference (*LD*) between the scores of standard and adaptive tree as:

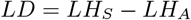

where *LH*_*S*_ is the LH of the standard tree and *LH*_*A*_ of the adaptive tree for a given MSA. The LD metric is measured in log-likelihood units (*LHUs*). Evidently, in cases where the adaptive tree has a higher score, the LD metric is negative. In 96.3% of empirical and 99.9% of simulated MSAs, the *LD* metric is below 2 LHUs. This means that, either the adaptive tree has a lower LH score but not by more than 2 LHUs, or has a higher LH score than the standard tree. Figure 1 in the Supplementary material summarizes the LD metrics over all empircal and simulated MSAs.

We further define the *relative LH difference (RLD)* as:

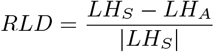

Again, in cases where the adaptive tree has a higher score, the RLD is negative. The results from our experiments indicate that:

- For 98.9% of the empirical datasets (9, 408 out of 9, 515 MSAs), the RLD is below 10^*−*3^. In the remaining 1.1%, the RLD does not exceed 0.02.
- For 99.9% of the simulated datasets (4, 997 out of 5, 000 MSAs), the RLD is below 10^*−*3^. In the remaining 3 datasets, the RLD does not exceed 0.01.

We can therefore claim that in 99% of all cases, adaptive RAxML-NG performs similarly, or even better, than the standard version in terms of log likelihood score. The RLD provides an intuition about the proximity of the two scores, even in cases where the absolute LH difference appears to be high. For example, in empirical dataset 11762_4 we observe the highest log-likelihood difference of 5, 237.10047 LHU among all MSAs for the standard-adaptive tree pair. The RLD metric, however, is only 0.003.

### 4.2 Significance tests and topological similarity

Figure 5 summarizes the results of the IQ-TREE 2 significance tests, conducted on all standard-adaptive tree pairs. The best scoring tree of each pair serves as the reference tree, and the hypothesis tested is whether the second tree is significantly worse under *any* of the statistical significance tests available in IQ-TREE 2. In approximately 94% of the cases, the two trees are statistically indistinguishable, while in 0.84% of the cases, adaptive RAxML-NG infers significantly better trees.

**Fig. 5.**
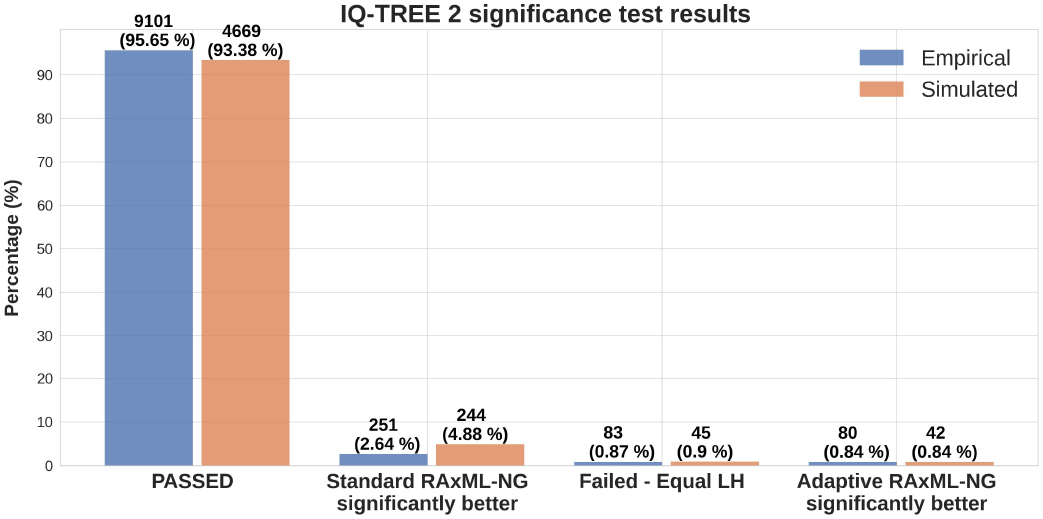
Results of IQ-TREE 2 significance tests. For 9 5% of the datasets, the inferred standard and adaptive trees are statistically indistinguishable. For approximately 1% of the datasets, adaptive RAxML-NG infers significantly better trees.

Figure 6 illustrates the relative RF distances between the two best trees found in all standard-adaptive tree pairs. We observe that, for datasets with a difficulty score below 0.5, the majority of pairs exhibits a relative RF distance below 0.2. This implies that the standard and adaptive version of RAxML-NG infer topologically similar trees on datasets with sufficient phylogenetic signal.

**Fig. 6.**
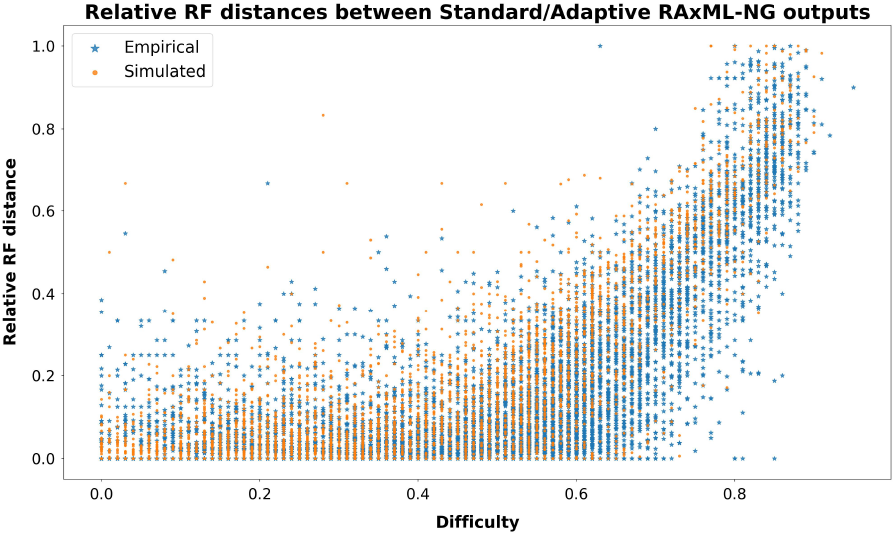
Relative RF distances between the two trees in all standard-adaptive tree pairs.

### 4.3 Speedups

For each dataset, we calculate the speedup by dividing the execution time of standard RAxML-NG by the execution time of adaptive RAxML-NG. Figure 7 summarizes the speedup distributions for empirical and simulated datasets over ten difficulty intervals. As expected, the results indicate substantial speedups (exceeding 5x) on easy and difficult datasets, since the number of independent tree searches performed by adaptive RAxML-NG is lower for these difficulty intervals.

**Fig. 7.**
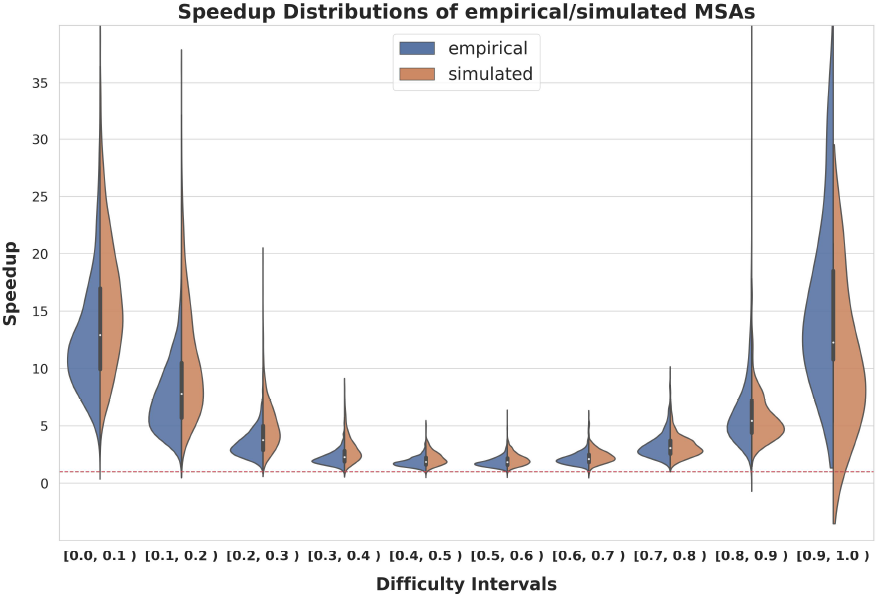
Speedup distributions for empirical/simulated datasets over 10 difficulty intervals. The red dashed line corresponds to a speedup of 1.

We further define the *Per-search Speedup (PS)* to be:

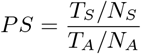

where *T*_*S*_, *T*_*A*_ are the overall execution times of standard and adaptive RAxML-NG, and *N*_*S*_, *N*_*A*_ is the number of independent tree searches conducted by each RAxML-NG version. In Table 1 we provide analogous statistics, the average value and the standard deviation, for the overall and per-search speedup distribution over 10 difficulty intervals.

**Table 1.**
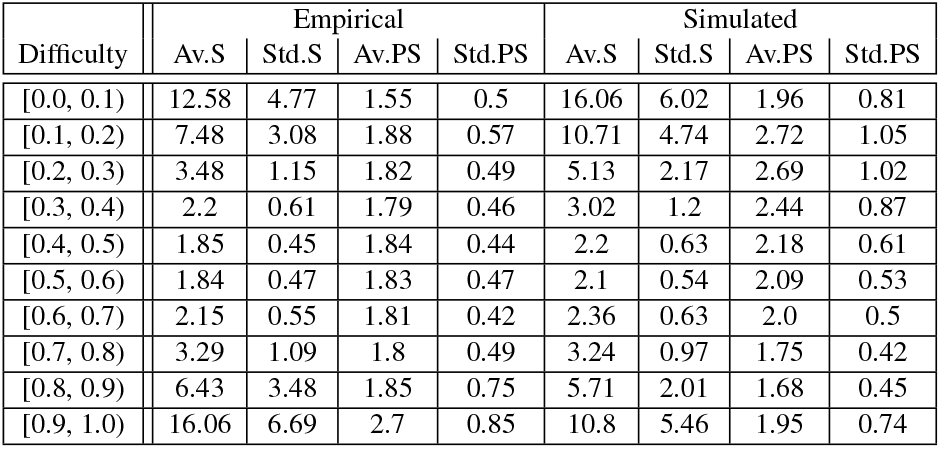
Average value and standard deviation of speedups and per-search speedups over 10 difficulty intervals. Av.S: average speedup, Std.S: standard deviation of speedups, Av.PS: average Per-search speedup, Std.PS: standard deviation of Per-search speedup

Finally, we calculate the overall run time for standard RAxML-NG by summing over the execution times of all standard RAxML-NG invocations. We calculate the overall run time for adaptive RAxML-NG analogously. We define the accumulated speedup to be the ratio of the overall accumulated execution times of the two versions. The accumulated speedup on empirical data is 3.11x and on simulated data is 4.27x.

## 5 CONCLUSIONS AND FUTURE WORK

We designed, implemented, and tested an adaptive version of RAxML-NG. We imported Pythia into the tree inference pipeline and modified the number of independent tree searches, as well as the thoroughness of the search heuristic, based on the predicted difficulty for the input MSA. For the vast majority of MSAs, our adaptive version performs equally well as the standard RAxML-NG version with respect to tree inference accuracy. As expected, the lower the difficulty score of the dataset, the higher the topological similarity between the two ML trees inferred from the standard and adaptive versions is. We achieve substantial overall and per-search speedups in our adaptive version, in particular on easy as well as difficult MSAs.

By introducing Pythia into the phylogenetic inference pipeline, we provide users an *a priori* estimate of the expected robustness of the final result, since the difficulty score directly reflects the amount of phylogenetic signal in the input MSA. The benefits of analyzing easy MSAs with our adaptive version are both the substantial speedups and the robustness of the final result.

Regarding future work, our first aim will be to efficiently parallelize the adaptive RAxML-NG version, by deploying a fine-grained parallelization scheme for the first ML inference on the first starting tree and either coarse-grained or automatic parallelization for the subsequent inferences on the remaining starting trees. The idea is to utilize more computational resources for the first ML inference such as to quickly establish a reference ML score and determine the 1% likelihood convergence interval, which will be used by the subsequent inferences for terminating the first stage early. Next, we intend to implement checkpointing in various phases of the tree search, to yield adaptive RAxML-NG more user-friendly. Lastly, we intend to test alternative heuristic tree search strategies and design improved heuristics for adaptive RAxML-NG to further improve likelihood scores and reduce runtime.

## ACKNOWLEDGMENT

This work was financially supported by the Klaus Tschira Foundation and by the European Union (EU) under Grant Agreement No 101087081 (Comp-Biodiv-GR).

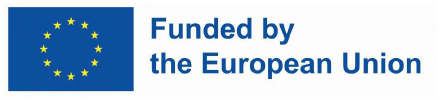

## Footnotes

1 Initially, Pythia used Random Forest Regression but this changed since its publication.

2 In this study, though, the comparison is between the updated versions of the tools: RAxML-NG, IQ-TREE 2, and FastTree 2.

3 By default RAxML-NG executes searches on 10 random and 10 MP starting trees

4 Two tree topologies are consecutive if they are separated by exactly one SPR and NNI round

5 This is not to be confused with the cases where the significance tests failed. We assign The label FAILED when the execution of IQ-TREE 2 was successful, but at least one of the tests failed to establish statistical connection

## Notes

https://github.com/togkousa/raxml-ng/tree/adaptive

https://cme.h-its.org/exelixis/material/raxml_adaptive_data.tar.gz

### Competing Interest Statement

The authors have declared no competing interest.

## REFERENCES

Bollback, J. P. (2002). Bayesian Model Adequacy and Choice in Phylogenetics. Molecular Biology and Evolution, 19(7), 1171–1180.

Felsenstein, J. (1981). Evolutionary trees from dna sequences: A maximum likelihood approach. Journal of Molecular Evolution, 17, 368–376.

Fitch, W. M. (1971). Toward Defining the Course of Evolution: Minimum Change for a Specific Tree Topology. Systematic Biology, 20(4), 406–416.

Guindon, S., Dufayard, J.-F., Lefort, V., Anisimova, M., Hordijk, W., and Gascuel, O. (2010). New Algorithms and Methods to Estimate Maximum-Likelihood Phylogenies: Assessing the Performance of PhyML 3.0. Systematic Biology, 59(3), 307–321.

Haag, J., Hübner, L., Kozlov, A. M., and Stamatakis, A. (2022a). The free lunch is not over yet – systematic exploration of numerical thresholds in phylogenetic inference. bioRxiv.

Haag, J., Höhler, D., Bettisworth, B., and Stamatakis, A. (2022b). From Easy to Hopeless—Predicting the Difficulty of Phylogenetic Analyses. Molecular Biology and Evolution, 39(12).p msac254.

Heath, L. S. and Ramakrishnan, N. (2010). Problem solving handbook in computational biology and bioinformatics. Springer-Verlag GmbH, 1st edition.

Höhler, D., Haag, J., Kozlov, A. M., and Stamatakis, A. (2022). A representative performance assessment of maximum likelihood based phylogenetic inference tools. bioRxiv.

Höhler, D., Pfeiffer, W., Ioannidis, V., Stockinger, H., and Stamatakis, A. (2021). RAxML Grove: an empirical phylogenetic tree database. Bioinformatics, 38(6), 1741–1742.

Kozlov, A. (2018). Models, Optimizations, and Tools for Large-Scale Phylogenetic Inference, Handling Sequence Uncertainty, and Taxonomic Validation. Ph.D. thesis, Karlsruhe Institute of Technology.

Kozlov, A. M., Darriba, D., Flouri, T., Morel, B., and Stamatakis, A. (2019). RAxML-NG: a fast, scalable and user-friendly tool for maximum likelihood phylogenetic inference. Bioinformatics, 35(21), 4453–4455.

Liu, K., Linder, C. R., and Warnow, T. (2011). Raxml and fasttree: Comparing two methods for large-scale maximum likelihood phylogeny estimation. PLOS ONE, 6(11), 1–11.

Mau, B., Newton, M. A., and Larget, B. (1999). Bayesian phylogenetic inference via markov chain monte carlo methods. Biometrics, 55(1), 1–12.

Minh, B. Q., Schmidt, H. A., Chernomor, O., Schrempf, D., Woodhams, M. D., von Haeseler, A., and Lanfear, R. (2020). IQ-TREE 2: New Models and Efficient Methods for Phylogenetic Inference in the Genomic Era. Molecular Biology and Evolution, 37(5), 1530–1534.

Morel, B., Barbera, P., Czech, L., Bettisworth, B., Hübner, L., Lutteropp, S., Serdari, D., Kostaki, E.-G., Mamais, I., Kozlov, A. M., Pavlidis, P., Paraskevis, D., and Stamatakis, A. (2020). Phylogenetic Analysis of SARS-CoV-2 Data Is Difficult. Molecular Biology and Evolution, 38(5), 1777–1791.

Morrison, D. A. (2007). Increasing the Efficiency of Searches for the Maximum Likelihood Tree in a Phylogenetic Analysis of up to 150 Nucleotide Sequences. Systematic Biology, 56(6), 988–1010.

Naser-Khdour, S., Minh, B. Q., Zhang, W., Stone, E. A., and Lanfear, R. (2019). The Prevalence and Impact of Model Violations in Phylogenetic Analysis. Genome Biology and Evolution, 11(12), 3341–3352.

Nguyen, L.-T., Schmidt, H. A., von Haeseler, A., and Minh, B. Q. (2014). IQ-TREE: A Fast and Effective Stochastic Algorithm for Estimating Maximum-Likelihood Phylogenies. Molecular Biology and Evolution, 32(1), 268–274.

Piel, W. H., Chan, L., Dominus, M. J., Ruan, J., Vos, R. A., and Tannen, V. (2009). TreeBASE v. 2: A Database of Phylogenetic Knowledge. e-BioSphere 2009.

Price, M. N., Dehal, P. S., and Arkin, A. P. (2010). Fasttree 2 – approximately maximum-likelihood trees for large alignments. PLOS ONE, 5(3), 1–10.

Robinson, D. and Foulds, L. (1981). Comparison of phylogenetic trees. Mathematical Biosciences, 53(1), 131–147.

Roch, S. (2006). A short proof that phylogenetic tree reconstruction by maximum likelihood is hard. IEEE/ACM Transactions on Computational Biology and Bioinformatics, 3(1), 92–94.

Rosenberg, M. S. and Kumar, S. (2001). Incomplete taxon sampling is not a problem for phylogenetic inference. Proceedings of the National Academy of Sciences, 98(19), 10751–10756.

Saitou, N. and Nei, M. (1987). The neighbor-joining method: a new method for reconstructing phylogenetic trees. Molecular Biology and Evolution, 4(4), 406–425.

St. John, K. (2016). Review Paper: The Shape of Phylogenetic Treespace. Systematic Biology, 66(1), e83–e94.

Stamatakis, A. (2011). Phylogenetic Search Algorithms for Maximum Likelihood, chapter 25, pages 547–577. John Wiley & Sons, Ltd.

Stamatakis, A. (2014). RAxML version 8: a tool for phylogenetic analysis and postanalysis of large phylogenies. Bioinformatics, 30(9), 1312–1313.

van Dorp, L., Acman, M., Richard, D., Shaw, L. P., Ford, C. E., Ormond, L., Owen, C. J., Pang, J., Tan, C. C., Boshier, F. A., Ortiz, A. T., and Balloux, F. (2020). Emergence of genomic diversity and recurrent mutations in sars-cov-2. Infection, Genetics and Evolution, 83, 104351.

Yang, Z. (2014). Molecular Evolution: A Statistical Approach. OUP Oxford.

Yang, Z. and Rannala, B. (1997). Bayesian phylogenetic inference using DNA sequences: a Markov Chain Monte Carlo Method. Molecular Biology and Evolution, 14(7), 717–724.

Zhou, X., Shen, X.-X., Hittinger, C. T., and Rokas, A. (2017). Evaluating Fast Maximum Likelihood-Based Phylogenetic Programs Using Empirical Phylogenomic Data Sets. Molecular Biology and Evolution, 35(2), 486–503.

