## Supplement for "Adaptive RAxML-NG: Accelerating Phylogenetic inference under Maximum Likelihood using dataset difficulty"

JOURNAL NAME

Vol. 00 no. 00 2023
Pages 1–4

### Supplement to "Adaptive RAxML-NG: Accelerating Phylogenetic inference under Maximum Likelihood using dataset difficulty"

Anastasis Togkousidis [1], Alexey M. Kozlov [1], Julia Haag [1], Dimitri Höhler [1], Alexandros Stamatakis [3,1,2]

[1]Computational Molecular Evolution Group, Heidelberg Institute for Theoretical Studies, Germany
[2]Institute of Theoretical Informatics, Karlsruhe Institute of Technology, Karlsruhe, Germany
[3]Biodiversity Computing Group, Institute of Computer Science, Foundation for Research and Technology - Hellas

Received on xx.xx.xxxx; revised on XXXXX; accepted on XXXXX

Associate Editor: xxxx xxx

#### 1 HEURISTIC

The pseudocode of our adaptive RAxML-NG heuristic is summarized in Algorithm 1. The functions are, by order of appearance:

- GETRANDSTARTINGTREES(): Takes as input the difficulty score of the MSA to be analyzed. It returns the number of random starting trees to be used by adaptive RAxML-NG, based on Figure 2a of the manuscript.
- GETPARSSTARTINGTREES(): Takes as input the difficulty score of the MSA to be analyzed. It returns the number of MP starting trees to be used by adaptive RAxML-NG, based on Figure 2a of the manuscript.
- OPTIMIZE(): This is the core function of the algorithm. It takes as input a random/MP starting tree, the difficulty score of the corresponding MSA, and a numerical parameter $\epsilon$. It conducts a full ML tree search based on the adaptive heuristic described in Section 3 of the manuscript and returns the ML tree and its log-likelihood score. The numerical parameter $\epsilon$ is used as a likelihood improvement threshold, that is, the tree search is terminated when the likelihood improvement does not exceed $\epsilon$.
- BLO(): Branch-length optimization routine. Takes as input a binary tree and optimizes its branches.
- MPO(): Model Parameter optimization routine. Takes as input a binary tree and optimizes its model parameters.
- NNI(): The NNI round routine. Takes as input a binary tree and applies a sequence of NNI moves, based on a greedy, hill-climbing heuristic. It returns the NNI-optimal tree.
- CONVERGED(): Takes as input the current log-likelihood score of best-scoring tree found so far during the tree search. It returns TRUE if the current score is less than $1\%$ worse than the reference score, otherwise it returns FALSE. As reference score we use the log-likelihood score of the best ML tree found so far during all (preceding) finished tree searches.
- FAST-SPR(): The Fast version of the SPR round. Takes as input a binary tree, applies a sequence of SPR moves based on a hill-climbing heuristic, and returns the optimized tree.
- SLOW-SPR(): The Slow version of the SPR round. Takes as input a binary tree, applies a sequence of SPR moves based on a hill-climbing heuristic, and returns the optimized tree.
- RFDIST(): The RF distance (Robinson and Foulds, 1981) function. Takes as input two binary trees and calculates the RF distance between them.
- GETSLOWSPRRADIUS(): Takes as input the difficulty score of the MSA being analyzed. It returns the Slow-SPR radius parameter calculated by adaptive RAxML-NG, based on Figure 2b of the manuscript.

#### 2 DIFFICULTY SCORE AND DATASETS

Figure 3 of the manuscript shows the distribution of empirical and simulated datasets over 10 difficulty intervals. We can observe that the proportion of datasets with a difficulty score exceeding 0.9 is comparatively low compared to the rest of the datasets. In the text we briefly mention that this phenomenon is associated with the definition of the difficulty score per se. Further, in Section 2.2 we provide a short description of the difficulty prediction paper (Haag *et al.*, 2022). In this work, members of our lab initially defined and calculated the inference difficulty of empirical MSAs by conducting 100 ML tree searches on each dataset using RAxML-NG. They provided the following definition of the difficulty score:

$$difficulty = \frac{1}{5} \cdot (RF_{all} + RF_{pl} + \frac{N^*_{all}}{N_{all}} + \frac{N^*_{pl}}{N_{pl}} + (1 - \frac{N_{pl}}{N_{all}}))$$

The five terms in the parenthesis, in the order they appear, are:

- $RF_{all}$ : The average relative RF distance between all pairs of trees in the 100 output ML trees.
- $RF_{pl}$ : The average relative RF distance between all pairs of trees in the plausible tree set.

© anonymous copyright 2023.

**Algorithm 1** Adaptive RAxML-NG heuristic

```
Input: MSA, difficulty, ϵ
Output: MLtree, MLnL
MLnL ← −∞, MLtree ← NONE                                  // Initialization
// Initialize random/MP trees and execute the heuristic
randTrees ← GetRandStartingTrees(difficulty)
parsTrees ← GetParsStartingTrees(difficulty)
for randTree in randTrees do                     // Execute for each random tree
    ( tmpMLtree, LnL ) ← Optimize(randTree, difficulty,ϵ)
    if LnL > MLnL then
        MLnL ← LnL, MLtree ← tmpMLtree
    end if
end for
for parsTree in parsTrees do                        // Execute for each MP tree
    ( tmpMLtree, LnL ) ← Optimize(parsTree, difficulty, ϵ)
    if LnL > MLnL then
        MLnL ← LnL, MLtree ← tmpMLtree
    end if
end for

function Optimize(tree, difficulty, ϵ)
    LnL ← −∞, impr ← TRUE                                 // Initialization
    // Initial BLO, MPO
    ( tree, LnL ) ← Blo (tree)
    ( tree, LnL ) ← Mpo (tree)
    // Easy and difficult datasets start with an NNI Round
    if difficulty < 0.3 OR difficulty > 0.7 then
        ( tree, LnL ) ← NNI (tree)
        ( tree, LnL ) ← Mpo (tree)
        if Converged(LnL) then go to SECOND STAGE
        end if
    end if

    // First stage, Fast-SPR + NNI
    sprRad ← 5, step ← 5, rf ← ∞, maxRad ← 25
    while NOT Converged(LnL) AND rf ! = 0 AND impr do
        ( newTree, newLnL ) ← Fast-SPR (tree, sprRad)
        ( newTree, newLnL ) ← NNI (tree)
        impr ← ( newLnL − LnL > ϵ )                              // Boolean
        rf ← RFdist ( tree, newTree)
        tree ← newTree, LnL ← newLnL
        if sprRad < maxRad then
            sprRad ← sprRad + step
        end if
    end while

SECOND STAGE:                                            // SLOW-SPR + NNI
    ( tree, LnL ) ← Mpo (tree)                          // Intermediate MPO
    impr ← TRUE
    sprRad ← GetSlowSPRradius(difficulty)
    while impr do
        ( tree, newLnL ) ← Slow-SPR (tree, sprRad)
        ( tree, newLnL ) ← NNI (tree)
        impr ← ( newLnL − LnL > ϵ ), LnL ← newLnL
    end while
    ( tree, LnL ) ← Mpo (tree)                                 // Final MPO
    return ( tree, LnL )                                // Return statement
end function

function Converged(LnL)
    if MLtree ! = NONE AND (MLnL−LnL)/|MLnL| < 0.01 then
        return TRUE
    end if
    return FALSE
end function

function GetSlowSPRradius(difficulty)
    if difficulty < 0.5 then
        return ⌊50· difficulty +5⌋
    else
        return ⌊−50· difficulty +55⌋
    end if
end function

function GetRandStartingTrees(difficulty)
    numTrees ← ⌊5.5·NormalDist(difficulty,0.5,0.2)⌋
    return min(numTrees, 10)                     // MIN(): minimum function
end function

function GetParsStartingTrees(difficulty)
    numTrees ← ⌊7.0·NormalDist(difficulty,0.5,0.25)⌋
    return min(numTrees, 10)                     // MIN(): minimum function
end function

function NormalDist(x,m,s)
    inv_sqrt_2pi ← 0.39894
    a ← (x - m) / s
    return inv_sqrt_2pi / (s · EXP(−0.5·a·a))   // EXP(): exponential function
end function
```

- $\frac{N^*_{all}}{N_{all}}$ : The number of unique tree topologies in the output ML tree set ($N^*_{all}$) divided by the total number of trees in the same set ($N_{all} = 100$).
- $\frac{N^*_{pl}}{N_{pl}}$ : The number of unique tree topologies in the plausible tree set ($N^*_{pl}$) divided by the total number trees in the same set ($N_{pl}$).
- $\frac{N_{pl}}{N_{all}}$ : The number of trees in the plausible tree set ($N_{pl}$) divided by the total number of trees in the output ML tree set ($N_{all} = 100$). We subtract this ratio from 1, leading to the full expression of the last term $(1 - \frac{N_{pl}}{N_{all}})$

Each term is a value between 0.0 and 1.0, leading to an average value between 0.0 and 1.0 that quantifies the overall difficulty. Since all terms are divided by 5, it follows that each term individually can contribute up to 0.2 units to the overall difficulty score. The reason why only a small proportion of MSAs have a difficulty score exceeding 0.9 is mainly associated with the last term in the parenthesis $(1-\frac{N_{pl}}{N_{all}})$. On easy datasets, almost all of the 100 output ML trees form part of the plausible tree set, since they share similar topologies and likelihood scores. Hence, the ratio $\frac{N_{pl}}{N_{all}}$ is close to 1 and the term $(1-\frac{N_{pl}}{N_{all}})$ is close to 0. The problem is that the majority of ML trees inferred from difficult MSAs are plausible as well. Although their topologies are contradicting, their likelihood scores are almost equal, and therefore the term $(1-\frac{N_{pl}}{N_{all}})$ is, again, close to 0. Even if the four remaining terms contribute their maximum value of 0.2 units each, the contribution of the very last term is, in most cases, lower than 0.1 (out of 0.2 units), summing up to an overall score which is lower than 0.9. Hence, only a small proportion of empirical MSAs in TreeBASE (Piel *et al.*, 2009) have a difficulty score exceeding 0.9. On the other hand, the remaining four terms in the parenthesis appear to alleviate this problem. On difficult MSAs, both the RF distance terms and the unique-topology ratios are close to 1, implying a high number of distinct and contradicting topologies in the output ML tree set. However, we believe that the definition of the difficulty score should be reformulated such that the difficult MSAs are distributed more uniformly within the difficulty score range $[0.7, 1]$.

Regarding the datasets used in our experiments, in Section 4 of the manuscript we describe in detail the filtering process which resulted in $9,515$ empirical and $5,000$ simulated MSAs in total. Out of the $9,515$ empirical MSAs, $8,052$ are unpartitioned DNA, $638$ are partitioned DNA, $817$ are unpartitioned amino-acid, and $8$ are partitioned amino-acid datasets. Further, from the $5,000$ simulated MSAs, $4,482$ are unpartitioned DNA, $18$ are partitioned DNA, $487$ are unpartitioned amino-acid, and $13$ are partitioned amino-acid datasets. We subsampled the simulated DNA datasets from the datasets used in study by Höhler *et al.* (2022). We simulated the AA MSAs based on a sample of RAxML Grove (Höhler *et al.*, 2021) datasets. RAxML Grove datasets contain files with inferred trees, their respective estimated substitution model parameters, as well as some statistical information about the analyzed MSA. In order to avoid too large simulated MSAs for the consecutive analyses, we selected datasets with an MSA number of unique sites (i.e., number of patterns) and number of taxa below the 95th percentile respectively. Then, we used the trees, respective substitution models (selected by the users of the RAxML web servers) and estimated model parameters to simulate MSAs using AliSim (Ly-Trong *et al.*, 2022). Overall, we used the following substitution models for AA MSA simulations: JTT+$\Gamma$ (Jones *et al.*, 1992)($29.8\%$), LG+$\Gamma$ (Le and Gascuel, 2008)($28.6\%$), Dayhoff+$\Gamma$ (Dayhoff, 1972)($26.7\%$), WAG+$\Gamma$ (Whelan and Goldman, 2001)($7,4\%$), Blosum62+$\Gamma$ (Henikoff and Henikoff, 1992)($4\%$), MtArt+$\Gamma$ (Abascal *et al.*, 2007)($2.7\%$), other ($0.8\%$).

For the analyses of all DNA MSAs we used the GTR+$\Gamma$ (Tavaré, 1986) model, and for analyses of the AA MSAs we used the LG model.

#### 3 COMMANDS

The user can invoke the adaptive RAxML-NG version using the `--adaptive` option when running standard RAxML-NG. The commands that we used to compare the two versions of RAxML-NG are the following:

Standard version:

```
./raxml-ng --threads 1 --msa {msa}
--model {model} --seed 0
```

Adaptive version:

```
./raxml-ng --adaptive --threads 1 --msa {msa}
--model {model} --seed 0
```

#### 4 ABSOLUTE LOG-LIKELIHOOD DIFFERENCES

Figure 1 summarizes the absolute log-likelihood differences ($LD$) for all standard-adaptive tree pairs, measured in log-likelihood units ($LHU$). We divide datasets into ten difficulty intervals. The heights of the bars correspond to the proportion of datasets, within the specified difficulty interval, where the LH difference lies within a specified LHU range. For example, in Figure 1a and on difficulty interval $[0, 0.1)$, there are $1,196$ datasets (purple bar) where the absolute LH difference of the standard-adaptive pair is between $0$ and $2$ LHU. This corresponds to approximately $72\%$ of the empirical datasets that have a difficulty score within $[0, 0.1)$.

#### ACKNOWLEDGMENT

This work was financially supported by the Klaus Tschira Foundation and by the European Union (EU) under Grant Agreement No 101087081 (Comp-Biodiv-GR).

Funded by the European Union

#### REFERENCES

Abascal, F., Posada, D., and Zardoya, R. (2007). Mtart: a new model of amino acid replacement for arthropoda. *Molecular biology and evolution*, **24**(1), 1–5.

Dayhoff, M. O. (1972). A model of evolutionary change in proteins. *Atlas of protein sequence and structure*, **5**, 89–99.

Haag, J., Höhler, D., Bettisworth, B., and Stamatakis, A. (2022). From Easy to Hopeless—Predicting the Difficulty of Phylogenetic Analyses. *Molecular Biology and Evolution*, **39**(12). msac254.

Henikoff, S. and Henikoff, J. G. (1992). Amino acid substitution matrices from protein blocks. *Proceedings of the National Academy of Sciences*, **89**(22), 10915–10919.

Höhler, D., Haag, J., Kozlov, A. M., and Stamatakis, A. (2022). A representative performance assessment of maximum likelihood based phylogenetic inference tools. *bioRxiv*.

Höhler, D., Pfeiffer, W., Ioannidis, V., Stockinger, H., and Stamatakis, A. (2021). RAxML Grove: an empirical phylogenetic tree database. *Bioinformatics*, **38**(6), 1741–1742.

Jones, D. T., Taylor, W. R., and Thornton, J. M. (1992). The rapid generation of mutation data matrices from protein sequences. *Bioinformatics*, **8**(3), 275–282.

Le, S. Q. and Gascuel, O. (2008). An improved general amino acid replacement matrix. *Molecular biology and evolution*, **25**(7), 1307–1320.

Ly-Trong, N., Naser-Khdour, S., Lanfear, R., and Minh, B. Q. (2022). AliSim: A Fast and Versatile Phylogenetic Sequence Simulator for the Genomic Era. *Molecular Biology and Evolution*, **39**(5). msac092.

Piel, W. H., Chan, L., Dominus, M. J., Ruan, J., Vos, R. A., and Tannen, V. (2009). TreeBASE v. 2: A Database of Phylogenetic Knowledge. *e-BioSphere 2009*.

Robinson, D. and Foulds, L. (1981). Comparison of phylogenetic trees. *Mathematical Biosciences*, **53**(1), 131–147.

Tavaré, S. (1986). Some probabilistic and statistical problems in the analysis of dna sequences. *Lect Math Life Sci (Am Math Soc)*, **17**, 57–86.

Whelan, S. and Goldman, N. (2001). A general empirical model of protein evolution derived from multiple protein families using a maximum-likelihood approach. *Molecular biology and evolution*, **18**(5), 691–699.

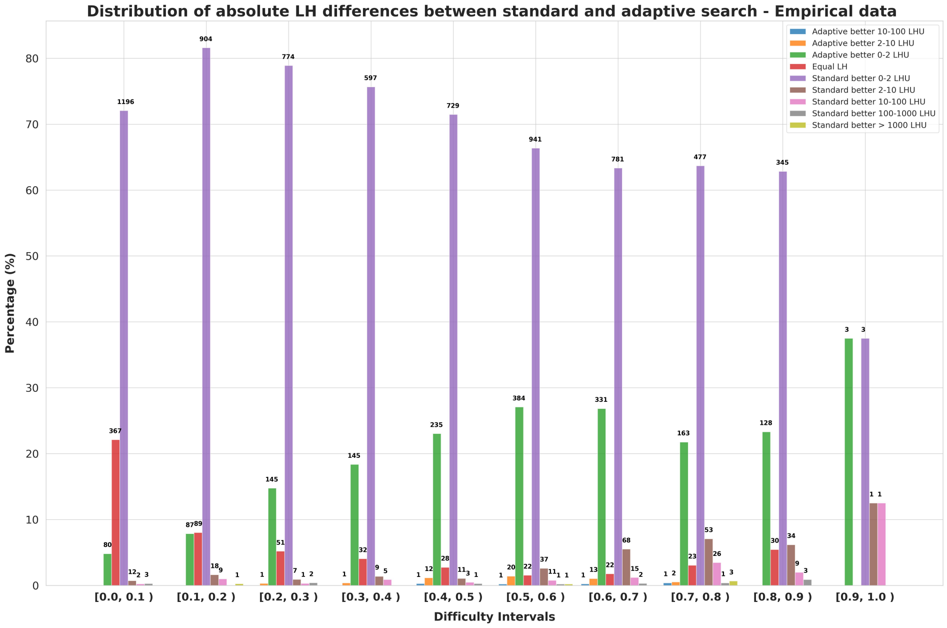


**(a)**

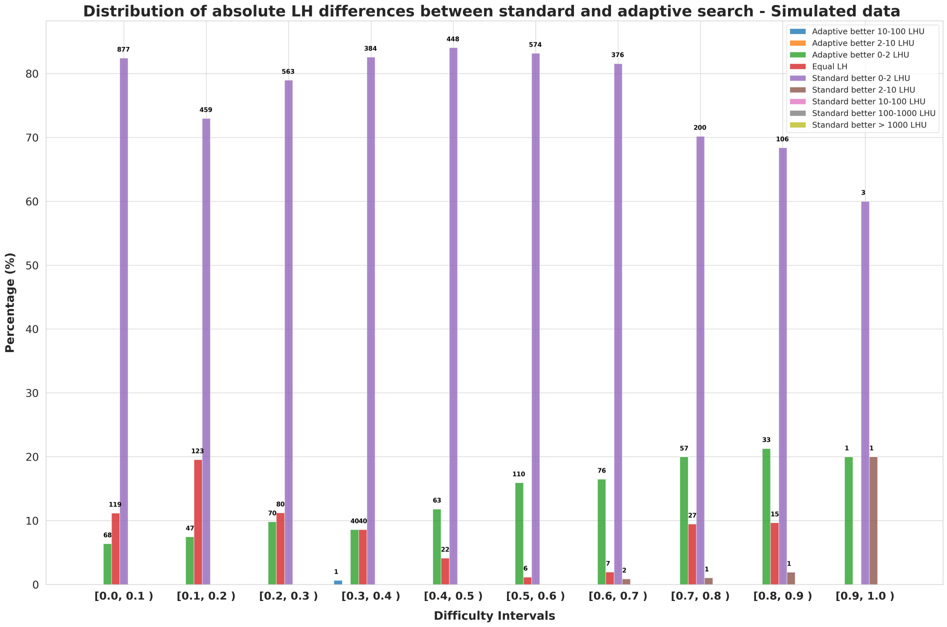


**(b)**

**Fig. 1.** Distributions of absolute log-likelihood differences in all standard-adaptive tree pairs, on **(a)** empirical and **(b)** simulated data. The abolute LH differences are measured in log-likelihood units ($LHU$). We divide the MSAs into ten difficulty intervals. The height of the bars corresponds to the proportion of datasets, within the specified difficulty interval, in which the score of standard and adaptive trees have an absolute difference within a range of LHU. The numbers at the top of the bars correspond to the number of datasets in this LHU range.
